# Fine Tuning of Histone Demethylase KDM6A/B Improves the Development of Nuclear Transfer Embryo

**DOI:** 10.1101/390484

**Authors:** Lei Yang, Lishuang Song, Xuefei Liu, Lige Bai, Guangpeng Li

## Abstract

Despite the success of the production of animals by somatic cell nuclear transfer (SCNT) in many species, the method is limited by a low efficiency. After zygotic genome activation (ZGA), a large number of endogenous retroviruses (ERVs) are expressed, including the murine endogenous retrovirus-L (MuERVL/MERVL). In this study, we generated a series of MERVL-reporter mouse strains to detect the ZGA event in embryos. We found that the majority of SCNT embryos exhibited ZGA failure, and histone H3 lysine 27 trimethylation (H3K27me3) prevented SCNT reprogramming. Overexpression of the H3K27me3-specific demethylase KDM6A, but not KDM6B, improved the efficiency of SCNT. Conversely, knockdown KDM6B not only facilitate ZGA, but also impede ectopic Xist expression in SCNT reprogramming. Furthermore, the knockdown of KDM6B increased the rate of SCNT-derived Duchenne muscular dystrophy embryonic stem cell establishment, indicate that these results not only provide insight into the mechanisms underlying failures of SCNT, but also may extend the applications of SCNT.

## Introduction

The metaphase II (MII) oocyte cytoplasm can reprogram somatic cell nuclei to the totipotent or pluripotent state via a series of sequential epigenetic events, including histone modifications, X chromosome reactivation, and pluripotency gene reactivation [1–4]. Somatic cell nuclear transfer (SCNT) has obvious advantages over other similar biotechnology techniques by enabling the generation of a new individual with an identical genome to that of the donor cell [5, 6]. However, SCNT-mediated reprogramming has a very low efficiency [7]. In particular, in mice, nearly half of SCNT embryos arrest at the pre-implantation stage and only 1–2% of SCNT embryos develop to term [8]. The molecular mechanisms underlying SCNT reprogramming are still unknown. Nevertheless, the successful reprogramming of human somatic cells by SCNT and the derivation of nuclear transfer embryonic stem cells (ntESCs) suggest that this is a promising approach [9–12].

A major feature of SCNT reprogramming is the global shift in gene expression from the somatic to the embryonic state. Zygotic genome activation (ZGA) occurs at the 2-cell stage in mice and at the 4-to 8-cell stage in pigs, bovines, and humans [13]. When the zygotic genome is first transcribed, a large number of retrotransposons are expressed, including endogenous retroviruses (ERVs), long interspersed nuclear elements, and non-autonomous short interspersed nuclear elements [14, 15]. MERVL repeats belong to type III ERVs and are specifically expressed at the 2-cell stage [16–21]. Hundreds of genes express chimeric transcripts with junctions to MERVL at the 5’ end, indicating that the long terminal repeats (LTRs) of MERVL serve as functional promoters [22, 23]. In the present study, we generated transgenic mouse lines containing a red fluorescent protein tandem dimeric tomato (tdTomato) reporter under the control of MERVL-LTR (MERVL::tdTomato). We used this unique reporting system to monitor ZGA in SCNT reconstructed embryos. Recent studies have indicated that ZGA in SCNT embryos is limited by histone H3 lysine 9 trimethylation (H3K9me3) barriers that preexist in the genome of donor cells [7, 24]. Previous studies have also indicated that treatment with pharmacological histone deacetylase and DNA methyltransferase inhibitors improves SCNT efficiency [25, 26]. However, SCNT efficiency is still not comparable to normal embryonic development, and it is likely that additional obstacles to SCNT reprogramming exist.

In this study, we demonstrated that ZGA failure is frequent in SCNT-generated embryos, and another prominent silencing marker, H3K27me3, is an obstacle for SCNT reprogramming. The overexpression of KDM6A, a H3K27me3-specific demethylase, facilitates ZGA-related gene expression in SCNT embryos. However, KDM6A-overexpressing SCNT embryos did not exhibit more efficient full-term development. On the contrary, KDM6B knockdown not only improved the blastocyst formation rate, but also increased the cloned embryo birth rate and ntES establishment efficiency. For future clinical applications of KDM6B knockdown-assisted SCNT, we derived blastocysts from DMD-deficient (X-chromosome linked muscular dystrophy, mdx) somatic cells and efficiently generated si6B-mdx-ntES. Thus, we established a highly efficient reprogramming method to improve SCNT for reproductive and therapeutic cloning.

## Results

### Most SCNT Embryos Exhibited ZGA and Developmental Failure

For the sensitive and convenient detection of ZGA events, we generated transgenic mouse lines containing a MERVL::tdTomato reporter (Fig 1A; Appendix Fig S1A). The cumulus cells from MERVL::tdTomato transgenic mice were used as nuclear donors for SCNT. As controls, intracytoplasmic sperm injection (ICSI) embryos were produced using the littermates of transgenic mice (Fig 1A; Appendix Fig S1B). As expected, the MERVL::tdTomato reporter was expressed at the late 2-cell stage (Fig 1B, C; Appendix Fig S1C, D; Movies EV1). We found that only 12% of SCNT embryos exhibited reactivation somatic MERVL::tdTomato at the 2-cell stage, while 92% of ICSI embryos exhibited reactivation (Fig 1E). MERVL encodes a canonical retroviral Gag protein [19]. We next verified the accuracy of the MERVL::tdTomato reporter by immunofluorescence (IF) and real-time quantitative PCR (qPCR), and these results are in accordance with the fluorescence images (Fig 1B, F; Appendix Fig S1E). To further confirm that the MERVL::tdTomato reporter can capture ZGA events, embryos were divided into tdTomato^+^ and tdTomato^−^ groups according to MERVL::tdTomato expression. The qPCR results showed that the expression levels of ZGA-related genes in tdTomato^+^ were significantly higher than those in the tdTomato^−^ group (Fig 1G; Appendix Fig S1F). After *in vitro* culture, for both ICSI or SCNT embryos, most tdTomato^+^ embryos developed to the blastocyst stage (97% and 89%, respectively). Surprisingly, we found that 18% SCNT-tdTomato^−^ embryos developed to the blastocyst stage, but none of the ICSI-tdTomato^−^ embryos reached the blastocyst stage, and most of them were blocked at the 2-cell stage (Fig 1H, I; Appendix Table S1). Notably, previous studies have shown that ZGA is essential for mouse embryonic development, as embryos will arrest at the 2-cell stage if ZGA is blocked [27]. Thus, MERVL::tdTomato could be used to monitor ZGA events in real time. Compared with ICSI embryos, a number of SCNT embryos arrested at various developmental stages (not limited to the 2-cell stage). Moreover, SCNT embryos are usually incapable of repressing some somatic genes inherited from donor cells [28, 29]. The expression of donor cell-specific genes in SCNT embryos could also lead to the development of a few SCNT-tdTomato^−^ embryos to blastocysts.

**Figure. 1.**
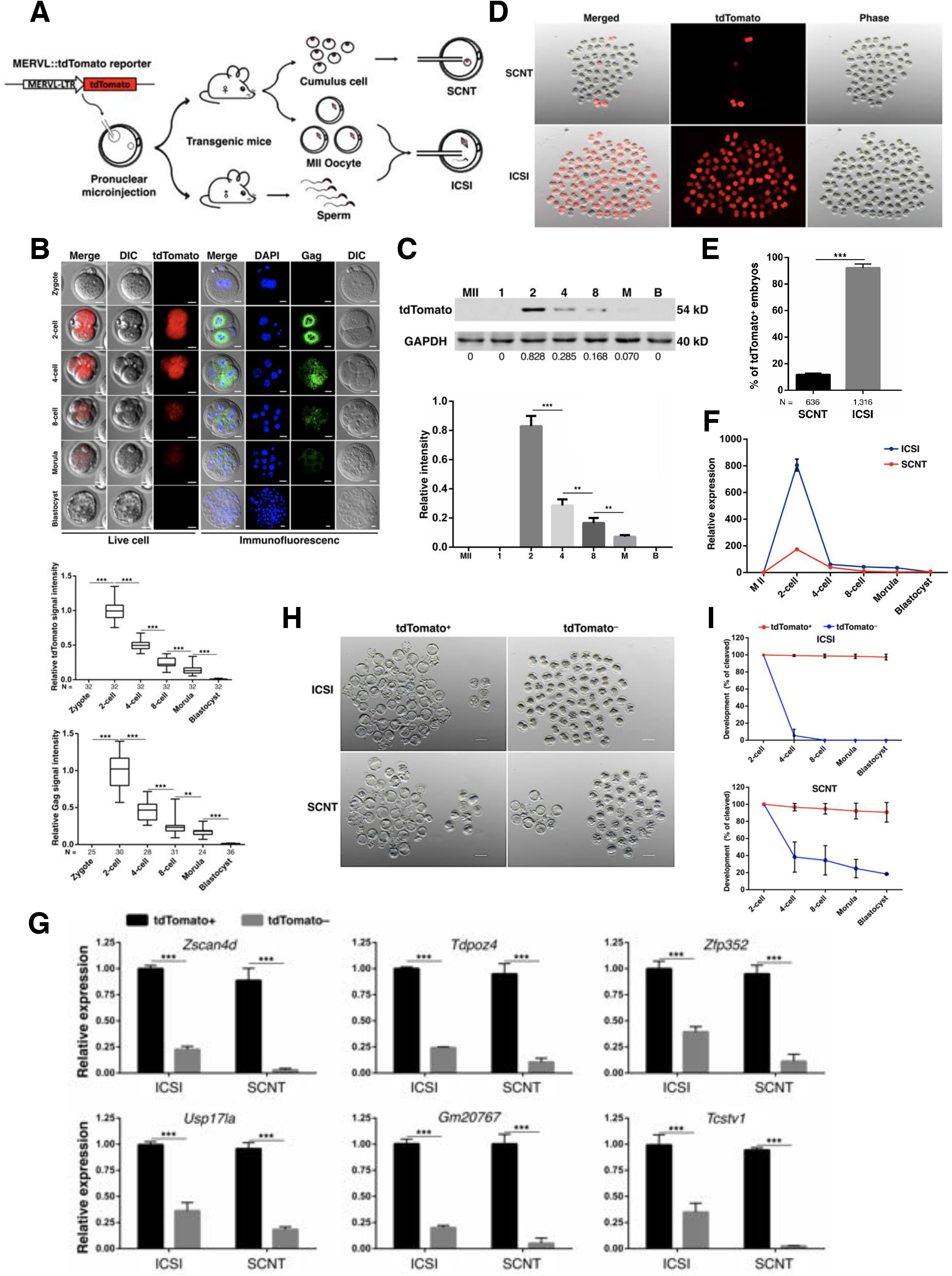
The most of SCNT reconstructed embryos are ZGA failure. A. Schematic view of the transgenic mice, ICSI and SCNT experiments. ♂ and ♀ indicated the male and female, respectively. B. Representative immunofluorescence and live-cell images of dynamics MERVL::tdTomato and Gag expression during embryos preimplantation development (upper). Quantification of tdTomato and Gag intensity (bottom). For the live-cell images, average intensity of tdTomato signal intensities relative to 2-cell stage embryos. For the immunofluorescence images, bar graphs showing the relative intensities of Gag/DAPI signal ratio. N, total number of embryos analyzed for each condition. Error bars, *s.d*., n ≥ 4. \*\**P* < 0.01, \*\*\**P* < 0.001 by two-tailed Student’s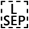*t*-test. Scale bar, 20 μm. C. Western blot analysis MERVL::tdTomato levels in MII oocyte and embryos at the indicated stages (upper). GAPDH was used as a loading control. Numbers below the western blots indicate band intensity (normalized to total GAPDH) measured by using ImageJ software. Quantification of western blot results (bottom). M, morula; B, blastocyst. Error bars, *s.d*., n = 3. \*\**P* < 0.01, \*\*\**P* < 0.001 by two-tailed Student’s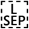*t*-test. Uncropped western blot and Ponceau S staining demonstrates equivalent loading of each lane are shown in Appendix figure S1D. D. Representative fluorescence image of 2-cell embryos derived from ICSI or SCNT. The SCNT embryos produced by transfer of MERVL::tdTomato cumulus cell into WT enucleated oocytes. ICSI embryos produced by MERVL::tdTomato sperm and MII oocytes from the littermates of transgenic mice. E. The summary of tdTomato^+^ 2-cell embryos derived from ICSI or SCNT. N, total number of embryos analyzed for each condition. Error bars, *s.d*., n ≥ 3. \*\*\**P* < 0.001 by two-tailed Studen’s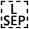*t*-test. F. The endogenous MERVL was up-regulated significantly in ICSI embryos compared with SCNT embryos at 2-cell stages, as determined by RT-qPCR. Error bars, *s.e.m*., n ≥ 3. G. RT-qPCR data for select ZGA genes activated following MERVL::tdTomato expression in mouse 2-cell embryos derived from ICSI or SCNT. Results were normalized based on the geometric mean of the expression levels of two reference genes (Ywhaz and Gapdh). Error bars, *s.e.m*., n = 3. \*\*\**P* < 0.001 by two-tailed Student’s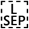*t*-test. H. Representative images of tdTomato^+^ and tdTomato^−^ embryos derived from ICSI or SCNT, after 4.5 days of culturing *in vitro*. Scale bar, 100 μm. I. Preimplantation development rates in the tdTomato^+^ and tdTomato^−^ embryos derived from ICSI or SCNT. The efficiency was calculated based on the number of 2-cell embryo that have been divided into tdTomato^+^ and tdTomato^−^ groups. Error bars, *s.d*., n ≥ 3.

### Effect of ZGA on SCNT Embryonic Development and ntESCs Derivation

Having established a correlation between MERVL::tdTomato and blastocyst formation, we next evaluated whether SCNT-tdTomato^−^ could develop to term. Because the IF assay requires fixation and/or denaturation, thereby preventing development, we used a live-cell imaging system to assess the full-term developmental ability of SCNT embryos (Fig 2A, B; Movies EV2). Based on tdTomato fluorescence, the SCNT blastocysts were grouped into SCNT-tdTomato^+^ and SCNT-tdTomato^−^. We detected fewer nuclei in SCNT-tdTomato^−^ blastocysts than in tdTomato^+^ blastocysts (Fig 2C, D). To further evaluate developmental ability *in vivo*, the SCNT-tdTomato^+^ and SCNT-tdTomato^−^ blastocysts with normal morphologies were used for embryo transfer. At embryonic day E6.5, no difference was observed between the SCNT-tdTomato^+^ and SCNT-tdTomato^−^ blastocysts in the implantation rate, as determined by the embryo retrieval rate (Appendix Fig S2A, B). However, 84% (16/19) of fetuses retrieved from tdTomato^+^ blastocysts had the typical morphology, with distinct embryonic and extraembryonic compartments, while none of the tdTomato^−^ fetuses were normal (0/29; Fig 2F). Furthermore, with respect to the ntESCs derivation efficiency, SCNT-tdTomato^+^ blastocysts had higher rates of attachment and ES establishment than those of SCNT-tdTomato^−^ blastocysts (Fig 2G, H; Appendix Fig S2C, D). Previous studies have demonstrated that Nanog is expressed in the inner cell mass (ICM) of blastocysts, and Cdx2 is expressed during formation of the blastocyst trophectoderm (TE), which represents the first step in embryo differentiation [30]. To gain further insights into blastocyst lineage segregation, the blastocysts derived from SCNT were subjected to IF staining of Nanog and Cdx2 (Fig 1E). In the SCNT-tdTomato^+^ blastocysts, Nanog and Cdx2 were exclusively localized to the nuclei of the ICM and TE, as previously reported in normal embryos [31]. By contrast, the Nanog and Cdx2 were localized to the cytoplasm of the ICM and TE in the SCNT derived tdTomato^−^ blastocysts. Thus, the Nanog and Cdx2 in SCNT-tdTomato^−^ embryos are mislocalization in a spatial manner, which may partially explain the developmental defects of SCNT-tdTomato^−^ embryos. We next examined the expression of somatic genes that have been reported to inhibit SCNT reprogramming [28, 32–35]; Fig 2I; Appendix Fig S2E). We found significant suppression of the expression of somatic cell genes at the 2-cell stage in the SCNT-tdTomato^+^ group, suggesting that these embryos have a greater degree of reprogramming than that of SCNT-tdTomato^−^ embryos.

**Figure. 2.**
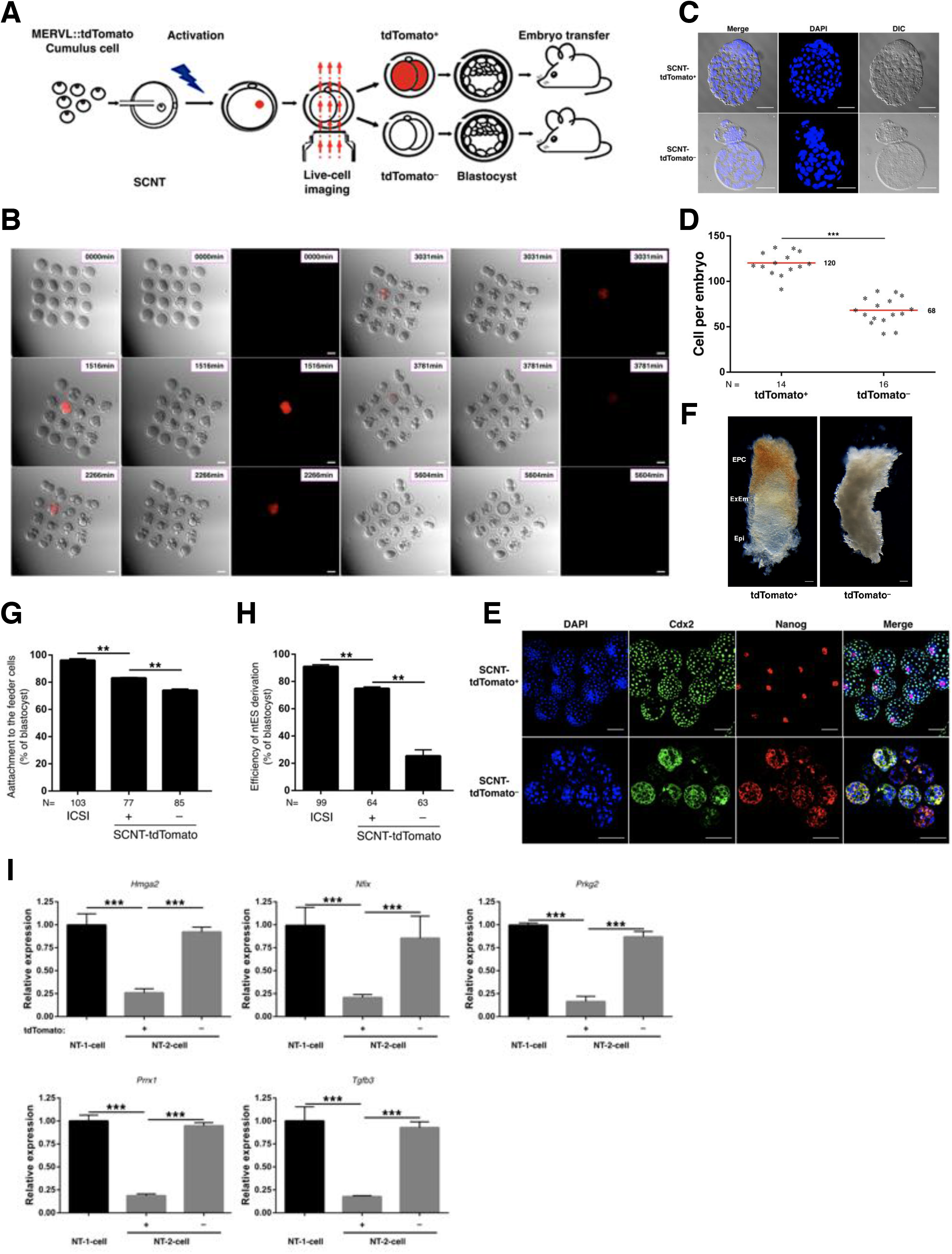
The effect of ZGA on the SCNT embryo quality. A. Schematics of the live-cell imaging experiments. embryo imaged from pronuclei until blastocyst stage and transferred to pseudopregnant females. B. Representative live-cell images of dynamics MERVL::tdTomato expression during SCNT embryos preimplantation development. The selected images from a series acquired every 15 min. The time after starting observation is shown on the upper right corner of each image. Scale bar, 50 μm. C. Representative DAPI staining of blastocysts of tdTomato^+^ and tdTomato^−^ SCNT embryos after 115 hr of culture *in vitro*. Scale bar, 50 μm. D. The tdTomato^+^ and tdTomato^−^ SCNT blastocyst cell numbers were determined by counting the DAPI-stained cells. N, total number of embryos analyzed for each condition. Red bars indicated the mean value. \*\*\**P* < 0.001 by two-tailed Student’s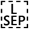*t*-test. E. Immunofluorescence images of tdTomato^+^ and tdTomato^−^ blastocysts derived from SCNT. Nanog (ICM) and Cdx2 (TE) were used as lineage markers. Representative images from ≥ 55 embryos analyzed in four independent micromanipulations are shown. Scale bar, 50 μm. F. Representative image of tdTomato^+^ and tdTomato^−^ SCNT embryos retrieved at E6.5. The tdTomato^+^ SCNT embryo displays normal egg cylinder morphology. By contrast, the tdTomato^−^ SCNT embryo shows abnormal morphology. Epi, embryonic epiblast; ExEm, extraembryonic ectoderm; EPC, ectoplacental cone. Scale bar, 50 μm. G. The bar chart showing the efficiency of attachment to the feeder cells of SCNT blastocysts. The efficiency was calculated based on the total number of blastocysts used for NTES derivation. N, total number of embryos analyzed for each condition. Error bars, *s.d*., n ≥ 3. \*\**P* < 0.01 by two-tailed Student’s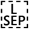*t*-test. H. The bar chart shows the efficiency of ntES derivation. The efficiency was calculated based on the total number of attached blastocysts for NTES derivation. N, total number of embryos analyzed for each condition. Error bars, *s.d*., n ≥ 3. \*\**P* < 0.01 by two-tailed Student’s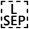*t*-test. I. RT-qPCR analysis for somatic cell genes in SCNT 1-cell embryos, 2-cell tdTomato^+^, and 2-cell tdTomato^−^ embryos. Results were normalized based on the geometric mean of the expression levels of two reference genes (Ywhaz and Gapdh). Error bars, *s.e.m*., n = 3. \*\*\**P* < 0.001 by two-tailed Student’s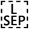*t*-test.

### Aberrant Reprogramming of H3K27me3 in the SCNT Embryos at the 2-cell Stage

In the SCNT mouse embryos, abnormalities in gene expression have been observed at the 2-cell stage, which corresponds to ZGA events. Furthermore, the epigenetic reprogramming of the somatic cell genome has been suggested as a key event in SCNT. We next determined the difference in epigenetic modifications between the SCNT and ICSI embryos at the 2-cell stage. Because histone H3K9me3 and H3K27me3 are correlated with gene silencing, while histone H3K4me3 leads to the initiation of gene transcription. The H3K4me3, H3K9me3, and H3K27me3 modifications of both ICSI and SCNT embryos were investigated (Fig 3A, B; Appendix Fig S3A, B, C). The IF assay indicated that H3K4me3 and H3K9me3 did not markedly differ between ICSI and SCNT-tdTomato^+^/tdTomato^−^ 2-cell embryos. In contrast, we found that H3K27me3 was specifically enriched in SCNT-tdTomato^−^ embryos, but moderate stain in SCNT-tdTomato^+^ and ICSI embryos. Contrary to H3K27me3 modification, the H3K27me2 did not differ between ICSI and SCNT derived embryos (Appendix Fig S3D). To further consolidate the IF results, we compared the H3K27me3 between different type embryos by Western-blot (WB). In the first set of experiments, SCNT-tdTomato^+^, SCNT-tdTomato^−^ and ICS-embryos were collected at 2-cell stage, the numbers of the embryos harvested for WB are 500, respectively. Furthermore, the polar bodies were also removed to avoid histone contamination. As IF results, in the short-exposure condition, H3K27me3 modification was effectively detected in the in the SCNT-tdTomato^−^ and cumulus cell (Fig. 3C; Appendix Fig S3E). When the embryos used for the WB were increased to 1,000 and under long-exposure condition, a weak band against theH3K27me3 was detected in the SCNT-tdTomato^+^ and ICSI samples (Fig. 3C; Appendix Fig S3E). Therefore, the H3K27me3 modification in the SCNT-tdTomato^+^ and ICSI 2-cell embryos is present at very low levels, but it can be detected. In addition, irrespective of whether female cumulus cells, male Sertoli cells, or mouse embryonic fibroblasts (MEFs) were used, the difference in H3K27me3 staining between the two types of SCNT embryos was also observed (Appendix Fig S3F). It is well known that fertilization unites two highly specialized haploid genomes with markedly different chromatin modifications within a single cell to form a diploid zygote. In the short period of the 1-cell stage, the two haploid genomes undergo dramatic asymmetric chromatin remodeling to reestablish transcriptional activation of zygotic gene expression [36]. We further investigated whether the difference in H3K27me3 modification also exists at the 1-cell zygote stage. We found that in 1-cell ICSI embryos, H3K27me3 signals were prominent in the maternal pronuclei, but not in the paternal pronuclei (Fig 3D; Appendix Fig S3G), which are consistent with previous study [37]. Unlike the asymmetric modifications in ICSI embryos, we detected strong H3K27me3 signals in all pseudo-pronuclei of SCNT embryos (Fig 3E). Furthermore, we also found that H3K27me3 levels were much higher in SCNT-tdTomato^−^ embryos than in SCNT-tdTomato^+^ or ICSI embryos at the morula stage (Fig 3F). According to the above results, we speculated that H3K27me3 is a natural key barrier preventing somatic cell nuclear reprogramming. We further examined the presence of H3K27me3 in bovine embryos, in which ZGA takes place during the 8-cell stage. As expected, the bovine intraspecific SCNT embryos also had much higher levels of H3K27me3 in the nuclei compared to those in the *in vitro* fertilization embryos at the 8-cell stage (Fig 3G, H). These results indicated that the H3K27me3 epigenetic barrier for SCNT-mediated reprogramming is shared across taxa.

**Figure. 3.**
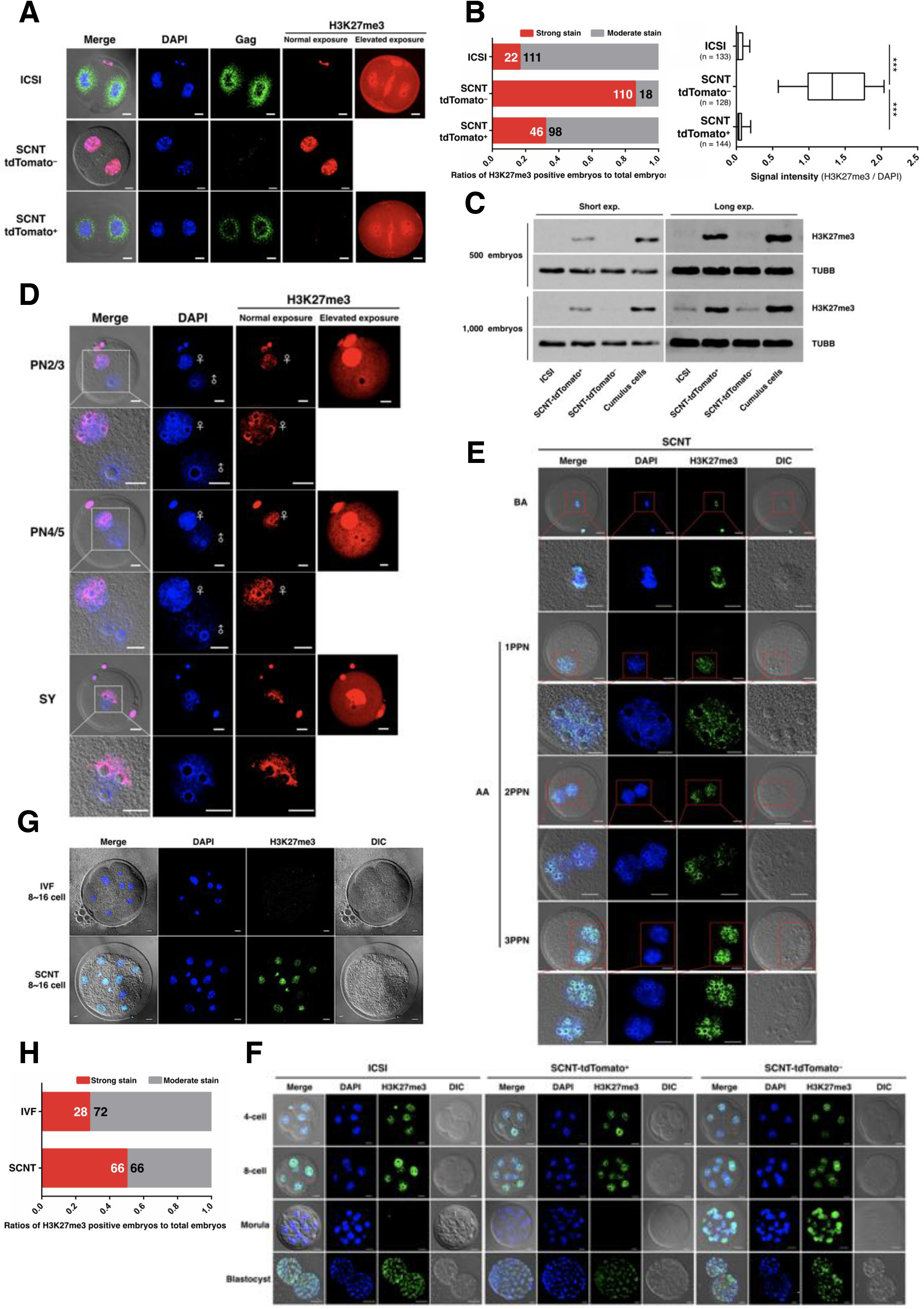
Abnormal H3K27me3 modification of SCNT embryos at the 2-cell stage. A. Immunofluorescence images of ICSI, SCNT-tdTomato^+^ and SCNT-tdTomato^−^ embryos. Embryos approximately 26hr after activation were fixed and processed for immunostaining with antibodies to H3K27me3 and Gag protein. ICSI embryos produced by MERVL::tdTomato sperm from the littermates of male mice. The images are representative examples of the quantification shown in Fig 3B. Representative images from ≥ 128 embryos analyzed in five independent micromanipulations are shown. Scale bar, 20 μm. B. Percentage of 2-cell ICSI and SCNT embryos with strong H3K27me3 staining and moderate staining (left). Numbers of the total embryos analyzed from five independent micromanipulations are shown in the bars. Boxplots for relative intensities of H3K27me3 from five independent experiments for each expression as in Fig 3A (right). Error bars, *s.e.m*., \*\*\**P* < 0.001 by two-tailed Student’s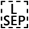t-test. C. Western blot results showing H3K27me3 levels in SCNT and ICSI embryos. α-Tubulin (TUBB) was blotted as a loading control. Uncropped western blot and Ponceau S staining demonstrates equivalent loading of each lane are shown in Appendix figure S3E. D. Representative immunofluorescent staining of H3K27me3 in ICSI embryos at the 1-cell stage. ICSI 1-cell embryos are characterised by the visualisation of two distinct pronuclei (2PN) 6 hours after sperm injection. H3K27me3 enrichment in the female-PN. The identity of the pronuclei was determined by their size and position relative to the polar body. Representative images from ≥ 95 embryos analyzed in four independent micromanipulations are shown. The optical Z-section series images are shown in Appendix figure S3G. ♂ and ♀ indicated the male and female, respectively. SY, syngamy. Scale bar, 20 μm. E. Representative immunofluorescent staining of H3K27me3 in SCNT embryos at the 1-cell stage. H3K27me3 enrichment in the SCNT embryos were observed with either 1 pseudo-pronucleus (1PPN), bipseudo-pronucleus (2PPN), or tripseudo-pronucleus (3PPN). BA, before activation; AA, after activation. Representative images from ≥ 55 embryos analyzed in three independent micromanipulations are shown. Scale bar, 20 μm. F. Dynamic appearance of H3K27me3 during early preimplantation development. Shown are representative images of embryos stained with DNA and H3K27me3. Negative staining of H3K27me3 could be observed in ICSI and SCNT-tdTomato^−^ embryo at morula stage. Representative images from ≥ 83 embryos analyzed in four independent micromanipulations are shown. Scale bar, 20 μm. G. Representative images of H3K27me3 immunostainings on bovine intraspecies SCNT embryos. IVF-derived zygotes were used as a control for comparison. These images are representative examples of the quantification shown in H. Representative images from ≥ 100 embryos analyzed in four independent micromanipulations are shown; Scale bar, 20 μm. H. The bar chart shows the percentages of strong and moderate H3K27me3 modifications of bovine embryos. Numbers of the total embryos analyzed from four independent micromanipulations are shown in the bars.

### Overexpression of KDM6A, but not KDM6B, Improves Preimplantation Development in SCNT Embryos

Having established that H3K27me3 is a barrier to somatic cell reprogramming, we next evaluated whether the removal of H3K27me3 could facilitate ZGA in SCNT embryos. We compared the expression levels of KDM6A and KDM6B, which are H3K27me3-specific demethylases, between ICSI embryos and SCNT embryos by RT-qPCR (Fig 4A, B). Neither KDM6A nor KDM6B was adequately activated in SCNT embryos. In addition, the expression levels of other KDMs in SCNT embryos were also lower than those in ICSI embryo (Appendix Fig S4A). To correct the H3K27me3 modification, the *in vitro* transcription vectors KDM6A and KDM6B tagged C-terminally with the hemagglutinin epitope (KDM6A-HA and KDM6B-HA) were constructed (Fig 4C). The exogenous HA ectopic expression vectors allowed us to track the KDM6A and KDM6B proteins in early embryos, without the use of specific antibodies. Strikingly, IF staining showed that ectopic expression of KDM6A or KDM6B markedly reduced the levels of H3K27me3 (Fig 4D; Appendix Fig S4D). Furthermore, other lysine methylation marks, including H3K9me3 and H3K4me3, were not affected (Appendix Fig S4B). Subsequently, we determined whether both KDM6A and KDM6B can improve the efficiency of SCNT reprogramming. We first injected KDM6A mRNA into enucleated MII oocytes (Fig 4E), and found that the overexpression of KDM6A mRNA significantly increased developmental efficiency (as determined by the blastocyst formation rate; Fig 4F). Surprisingly, the efficiency of SCNT was greatly reduced by injecting KDM6B mRNA into enucleated MII oocytes prior to SCNT (even at low doses; Fig 4F; Appendix Fig S4C). We also noticed that a KDM6A concentration of 20 or 50 ng/μl substantially improved the SCNT blastocyst development rate, while concentrations of KDM6A mRNA over 200 ng/μl were detrimental to embryonic development (Fig 4G; Appendix Fig S4C). To further investigate whether KDM6A overexpression improved the efficiency of full-term development, we transferred the SCNT embryos derived above into surrogates. For most transfers, pregnancies were established and maintained until day E8.5 and the fetuses were retrieved on that day (Fig 4H left). We found that the embryo retrieval rate for the group injected with KDM6A mRNA was substantially greater than that of directly transferred SCNT embryos (Fig 4I). Unexpectedly, only implantation sites and degenerated embryos were observed on day E19.5, suggesting that KDM6A-treated SCNT fetuses failed and were reabsorbed at E8.5–19.5 (Fig 4H right). These results indicate that the overexpression of KDM6A (but not KDM6B) improved pre-implantation development, but could not improve the rate of full-term development in SCNT fetuses.

**Figure. 4.**
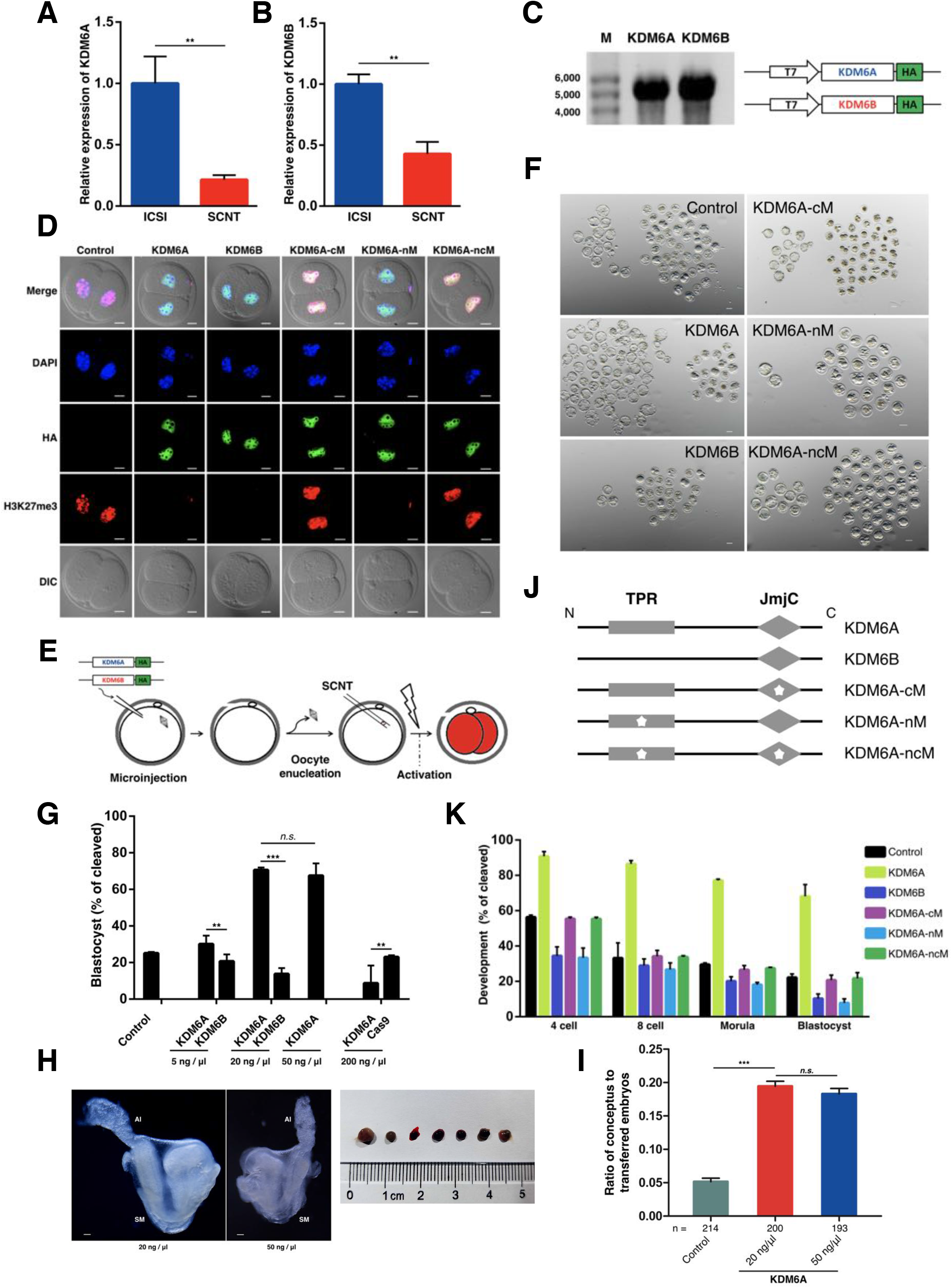
Overexpression of KDM6A only improves the blastocyst formation rate of SCNT embryos, but not full-term development. A, B RT-qPCR analysis of KDM6A (A) and KDM6B (B) mRNA levels in SCNT 2-cell embryo. Data shown are mean expression values relative to *Gapdh*. The value in ICSI control was set as 1. Error bars, *s.e.m*., n ≥ 3. \*\**P* < 0.01 by Student’s t-test. C. The sketch of KDM6A and KDM6B *in vitro* transcription vector (right), and the integrity of *in vitro* transcripted mRNA was confirmed by electrophoresis with formaldehyde gels (left). M, marker; T7, *in vitro* transcription promoter; HA, hemagglutinin epitope tag. D. Immunostaining of SCNT embryo for H3K27me3 and HA epitope tag after injection of different mRNA as indicated. Shown are representative images in three independent experiments. Representative images from ≥ 187 embryos analyzed in four independent micromanipulations for each condition are shown. Scale bar, 20 μm. E. Schematic illustration of mRNA injection into oocytes and SCNT. F. Representative images of SCNT embryos at 115 h after injection of different mRNA as indicated. Scale bar, 50 μm. G. The bar chart showing the efficiency of blastocyst formation. Injection of KDM6A mRNA improved the preimplantation development rate of SCNT embryos. Related to Appendix Figure S4C. Error bars, *s.e.m*., n ≥ 3. \*\**P* < 0.01, \*\*\**P* < 0.001 by two-tailed Student’s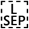*t*-test. *n.s*., not significant.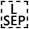 H. Phenotypic analysis of E8.5 SCNT mouse embryos injection with 20 ng/μl or 50 ng/μl KDM6A mRNA (left). Representative images of the KDM6A injected embryos at E19.5 (right). The injected SCNT embryos were only obtained degenerated embryos. AI, allantois; SM, somite. Scale bar, 50 μm. I. The retrieved rate of embryos 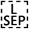assessed at E8.5. The numbers at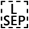the bottom of the bars indicated the total number of transferred embryos. *n.s*., not significant; \*\*\**P* < 0.001 by two-tailed Student’s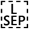*t*-test. J. KDM6A are broadly expressed proteins characterized by N-terminal TPRs and C-terminal JmjC domain. In KDM6B the only clearly identifiable domain is the C-terminal JmjC domain. Schematic diagram depicts the position of highly conserved TPR and JmjC domain, which are mutated to abolish the protein-protein interactions and demethylation functions of KDM6A. K. Preimplantation development rates in the KDM6A-HA, KDM6B-HA, or KDM6A-cM/-nM/-ncM-HA mRNA-injected and non-injected Control SCNT groups. The efficiency was calculated based on the number of cleavage embryo. Error bars, *s.d*., the total numbers of cleavage embryos in each condition (KDM6A-HA, KDM6B-HA, KDM6A-cM/-nM/-ncM-HA, and Control) from three independent experiments were 275, 199, 290, 221, 244, and 286, respectively.

Both KDM6A and KDM6B are Jumonji (JmjC) domain containing proteins and catalyze the removal of trimethylation from histone H3K27 by using a hydroxylation reaction with iron (Fe^2^+) and α-ketoglutarate (α-KG) as cofactors [38, 39]. The jumonji gene was named for a mutation in mice that causes abnormal cruciform neural grooves (in Japanese, jumonji means cruciform). As shown in Fig 4J, KDM6B shows high homology and structural relationship to KDM6A, especially in the JmjC domain, but lacks the tetratricopeptide (TPR) domain, which are assumed to mediate protein-protein interactions [40, 41]. In order to further compare the differences between KDM6A and KDM6B in SCNT reprogramming. We synthesized KDM6A-HA expression vectors with different loci mutation, and injecting different type mRNA into SCNT embryos (Fig 4J; Appendix Fig S4E). When KDM6A-cM-HA (JmjC domain mutant) or KDM6A-ncM-HA (TPR and JmjC double mutant) was ectopically expressed in SCNT embryos, no reduction in H3K27me3 methylation levels was observed (Fig 4D; Appendix Fig S4D), which demonstrating that the demethylation activity is dependent on JmjC domain. Furthermore, the blastocyst formation rate of SCNT embryos was greatly reduced when KDM6A-nM-HA was injected, which was similar to that of KDM6B injected SCNT embryos (Fig 4F, K; Appendix Table S2). Compared with the control group, the efficiency of SCNT was no different by injecting KDM6A-cM-HA or KDM6A-cnM-HA into SCNT embryos. These results suggesting that the TPR and JmjC domain were required for KDM6A rescue the poor developmental phenotype of SCNT embryos, and indirectly indicate that TPR domain may mediate protein-protein interactions for moderate KDM6A activity in the SCNT reprogramming.

### KDM6B Knockdown Increased the Expression of KDM6A and Blastocyst Formation Rate

As described above, the ectopic overexpression of KDM6A mRNA at low concentrations improved the SCNT efficiency. We have previously shown that mouse parthenogenetic embryos in which KDM6B is knocked down exhibited a moderate increase in KDM6A expression [42]. We speculated that KDM6B knockdown could facilitate ZGA and improve SCNT efficiency. To verify this hypothesis, we designed and constructed short interfering RNA (siRNA) specifically targeting KDM6A and KDM6B (Fig 5D; Appendix Fig S5A). A siRNA without any specificity to KDM6A/B or other genes was constructed as an siRNA-control. As expected, the qPCR results demonstrated that the decrease in KDM6A or KDM6B expression was accompanied by an increase in KDM6B or KDM6A expression, respectively (Fig 5A). Furthermore, a marked decrease H3K27me3 levels were observed when injected with either KDM6A or KDM6B siRNA (Fig. 5C; Appendix Fig S5C). The WB results also confirmed this phenomenon at another protein levels (Fig. 5B; Appendix Fig S5B). These findings suggest that KDM6A and KDM6B are functionally redundant and compensate for each other in SCNT embryos; that is interference of either KDM6A or KDM6B, the levels of the other will increase. At the beginning of the knockdown assay, we noticed that the pluripotency genes Oct4, Sox2, and Nanog are acquire the H3K27me3 mark as they get repressed during ESCs differentiation [43, 44]. In addition, KDM6B also regulate the Hox gene expression, which are essential for regulating cell differentiation and the formation of body structures during early embryonic development. In order to avoid injuries caused by knockdown KDM6B, we next tested a serial dilution of siRNA-6B to determine the knockdown efficiency (Fig 5D). Briefly, the optimal injection concentration of siRNA-6B in our experiment was 10 μM. We then injected siRNA-6B into recipient MII oocytes before SCNT (Fig 5E). We next wondered whether there were differences between KDM6B knockdown and KDM6A overexpression in the rate of SCNT blastocyst formation. Notably, the injection of KDM6B siRNA before SCNT increased the blastocyst rate to 70.8%, which did not differ significantly from the rate observed for KDM6A mRNA injection alone (70.3%; Fig 5F, G; Appendix Table S3). Furthermore, using Sertoli or MEF cells, the injection of siRNA-6B before SCNT also increased the blastocyst formation rate (Fig 5G; Appendix Fig S5D and Table S3). When we performed the same trials for bovine intraspecific SCNT, siRNA-6B injection also improved the developmental efficiency (Fig 5G; Appendix Fig S5D and Table S3). Interestingly, when we decrease the expression of KDM6A by injecting siRNA-6A into SCNT embryo, and found the blastocyst formation rate was significantly reduced (Fig 5F, G; Appendix Table S3). To further examine whether the positive effect of siRNA-6B on SCNT embryonic development is dependent on the observed increase KDM6A expression. We next double injection of siRNA-6A-6B into SCNT embryo. We observed significantly lower developmental potential for siRNA-6A-6B injected SCNT embryos, with the majority arresting at the 2-stage and only a few reaching the blastocyst stage (Fig 5F, G; Appendix Table S3). We also compared the ZGA related genes expression between different type siRNA injected SCNT embryos via RT-qPCR. Similarly, the qPCR results showed that the expression of ZGA related genes are decreased in SCNT embryo with siRNA-6A or siRNA-6A-6B injected compared with the control (Fig 5H). These results suggest that the overexpression of KDM6A or knockdown of KDM6B can improve the efficiency of SCNT reprogramming.

**Figure. 5.**
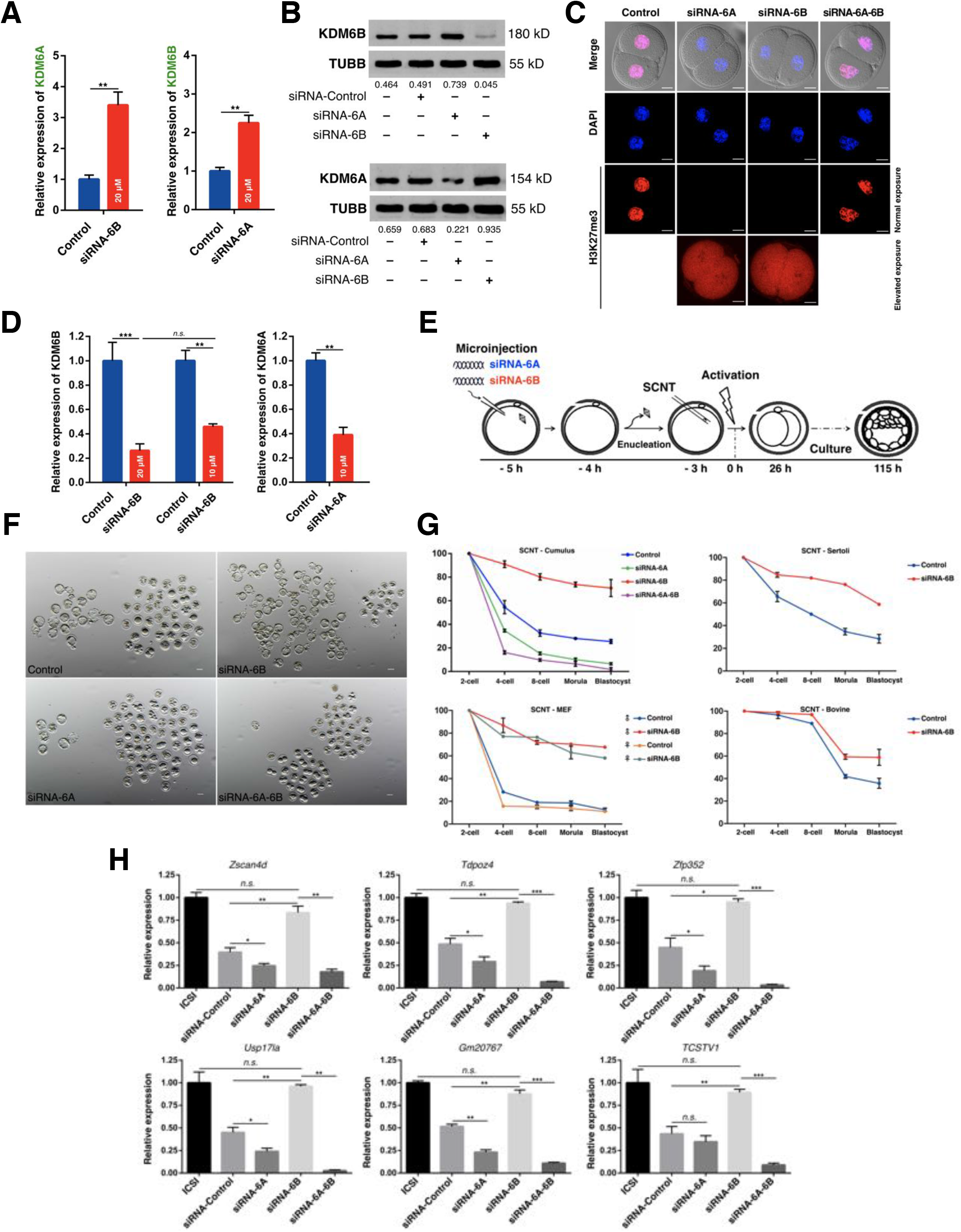
KDM6B knockdown 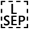greatly improved the preimplantation development rate of SCNT embryos. A. The bar chart shows the KDM6A/B transcript levels in SCNT embryos 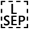with 20 μM siRNA-6A/B. All data are mean expression relative to *Gapdh* with control siRNA injected SCNT embryos normalized to 1.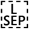Error bars, *s.e.m*., n = 3, \*\**P* < 0.01 by Student’s t-test. B. Western blot showing the expression of KDM6A and KDM6B in SCNT embryos after injection of different siRNA as indicated. Protein lysates from 1,500 embryos were loaded in each lane. The results of one representative of two independent experiments are presented. Uncropped western blot and Ponceau S staining demonstrates equivalent loading of each lane are shown in Appendix Fig S5B. C. Immunofluorescence staining results showing H3K27me3 of 2-stage SCNT embryos. Embryo was injected with siRNA as indicated. The H3K27me3 levels between control and double injected with siRNA-6A-6B cannot observe any difference, but a marked decrease was observed when injected with either siRNA-6A or siRNA-6B. scale bar, 20 μm. D. RT-qPCR analysis of KDM6A/B in SCNT embryos injected with siRNA-6A or siRNA-6B. The SCNT embryos were subject to injection of 10 μM or 20 μM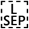siRNA as indicated. Data are mean expression relative to Gapdh with siRNA-control normalized to 1. Error bars, *s.e.m*., n = 3. \*\**P* < 0.01, \*\*\**P* < 0.001 according to two-tailed Student’s *t*-test. *n.s*., not significant. E. Schematic illustration of siRNA injection into oocytes and SCNT. F. Representative images of SCNT embryos at 115 h after injection of different siRNA as indicated. These images are representative examples of the quantification shown in G. n ≥ 3. Scale bar, 50 μm.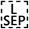 G. Injection of siRNA-6B improved the preimplantation development rate of SCNT embryos. Both cumulus cells, Sertoli cells, and C57 MEF cells were used as donor cells. The bovine intraspecies SCNT embryos derived from bovine ear fibroblast cells. Shown is the percentage of embryos that reached the indicated stages. ♂ and ♀ indicated the male and female, respectively. Error bars, *s.d*., n ≥ 3. H. RT-qPCR analysis for select ZGA genes in ICSI and SCNT embryos. The SCNT embryo was injected with siRNA as indicated. Results were normalized based on the geometric mean of the expression levels of two reference genes (Ywhaz and Gapdh). Error bars, *s.e.m*., n = 3. \**P* < 0.05, \*\**P* < 0.01, \*\*\**P* < 0.001 according to two-tailed Student’s *t*-test. *n.s*., not significant.

### KDM6B Knockdown Increased the SCNT Embryo Birth Rate as well as the Efficiency of DMD-specific ntES Derivation

Reactivation of pluripotency genes is a major event for the successful reprogramming of somatic cells to the blastocyst state. In particular, the transcription factor Oct3/4 (Pou5f1) is expressed in the ICM of the blastocyst stage, which is an effective indicator of embryonic quality. To determine the extent to which the injection of siRNA-6B could overcome ZGA defects in the SCNT embryos, we intercrossed MERVL::tdTomato with Oct4::EGFP transgenic mice (expressing the enhanced green fluorescence protein controlled by the Oct3/4 promoter) to produce MERVL::tdTomato/Oct4::EGFP dual reporter mice (Fig 6A; Appendix Fig S6A). Similar to other somatic cells, the cumulus cells and sperm did not express tdTomato and EGFP (Appendix Fig S6B). As expected, 40.7% of the siRNA-6B injected SCNT embryos exhibited tdTomato expression at the 2-cell stage, whereas only 3.5% of the siRNA-control group exhibited tdTomato fluorescence (Fig 6B, C). We also found weak, but substantial expression of Oct4::EGFP in the siRNA-6B-injected blastocysts (25/33, 75.6%), but not in the siRNA-control injected embryos (0/25; Fig 6D). We further compared the Oct4::EGFP mRNA levels between ICSI and siRNA-injected SCNT blastocysts. The qPCR results also showed that Oct4::EGFP mRNA expression was higher in SCNT blastocysts injected with siRNA-6B than in controls (Fig 6E). Moreover, the siRNA-6B-injected embryos contained a greater total cell number than that of the control blastocysts (118 in siRNA-6B injected *vs*. 67 in control; Appendix Fig S6C). We next investigate whether this positive effect could be contributing to cloned mice birth. For this purpose, siRNA-6B-injected SCNT embryos were transferred at the 2-cell stage into pseudo-pregnant females. Caesarian section at E19.5 revealed that the 6.0% (16/265; six twins) of transferred siRNA-6B-injected SCNT embryos developed to term, while none of the 120 transferred control embryos developed to term (Fig 6F; Appendix Fig S6D, E). To better characterize post-implantation development, we retrieved the siRNA-6B-injected SCNT conceptus at E15.5. The results showed that 43.5% (17/39) of the implantation sites still contained a fetus (nearly half of which were still alive; Appendix Fig S6F). Upon closer examination, we found one fetus (1/17) with intestinal fistula and skull closure defects (Appendix Fig S6D).

**Figure. 6.**
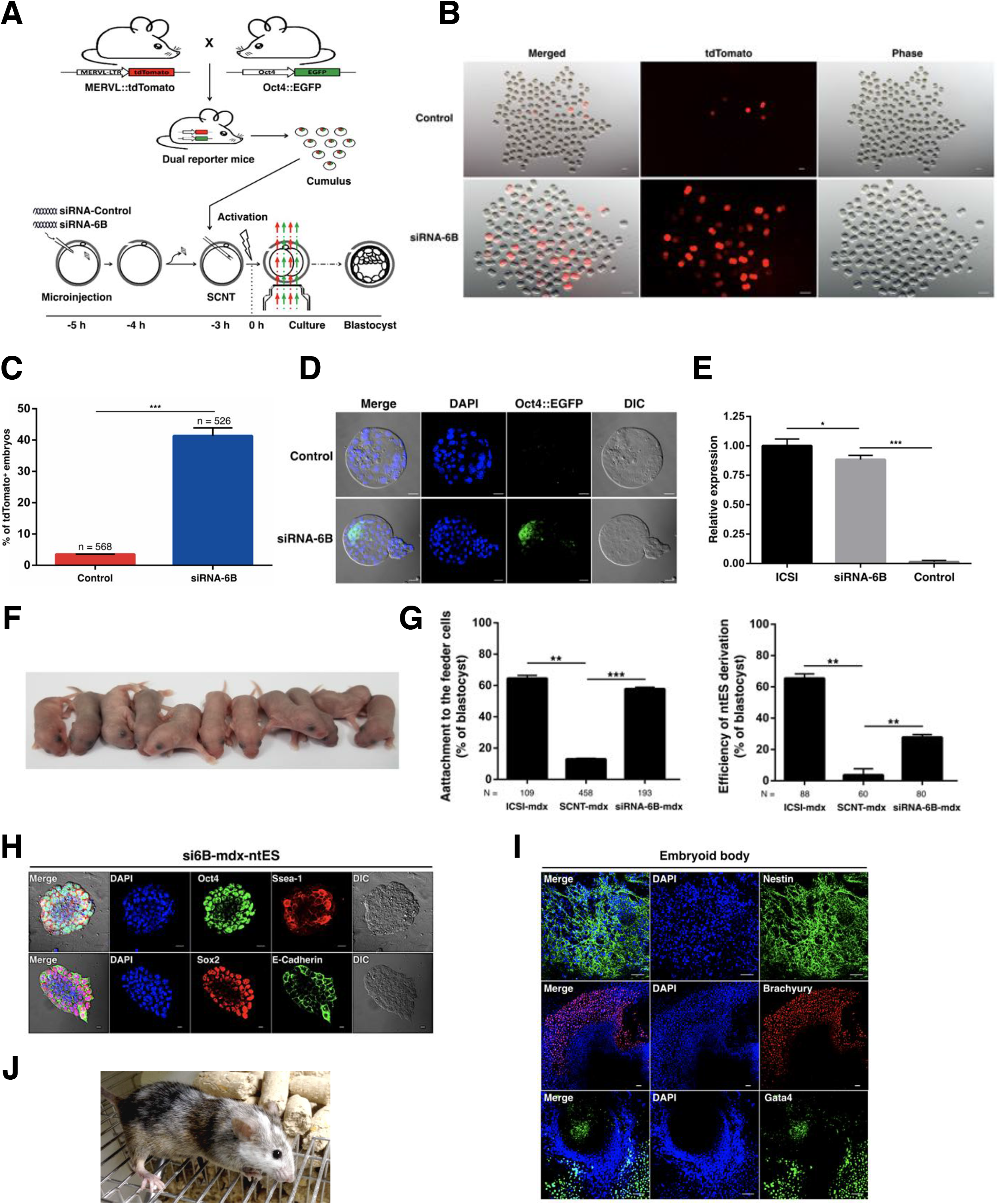
KDM6B knockdown improved the SCNT embryo birth rate and DMD-ntES derivation. A. Experimental scheme for generating dual reporter mice and SCNT. B. Representative fluorescence images of siRNA-control or siRNA-6B injected SCNT embryos. Scale bar, 50 μm. C. Quantification of embryos that expression tdTomato after injection with siRNA-control or siRNA-6B. Numbers of observed embryos are indicated. n ≥ 3, Error bars, *s.e.m*., \*\*\**P* < 0.001 according to two-tailed Student’s t-test. D. Representative images of green fluorescence in Oct4::EGFP reconstructed blastocysts derived from injected siRNA-control or siRNA-6B SCNT embryos. Representative images from ≥ 25 embryos analyzed in three independent micromanipulations are shown. Green fluorescence indicates that the Oct4::EGFP transgene has been expressed. Scale bar, 50 μm. E. Oct4::EGFP was up-regulated significantly in ICSI embryos compared with SCNT embryos at blastocyst stages, as determined by RT-qPCR. Data shown are mean expression values relative to *Gapdh*. The value in ICSI embryos was set as 1. Error bars, *s.e.m*., n ≥ 3. \**P* < 0.05, \*\*\**P* < 0.001 by two-tailed Student’s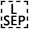*t*-test. F. Representative image of cloned mice derived by siRNA-6B injected SCNT embryos. G. Bar graph showing the efficiency of attachment to the feeder cells (left) and si6B-mdx-ntES derivation (right). The mdx sick mice tail-tip fibroblasts as nuclear donors. N, total number of embryos analyzed for each condition. H. Immunostaining images of si6B-mdx-ntES expressed pluripotency markers. Scale bar, 50 μm. I. The si6B-mdx-ntES possessed multiple-differentiation potential, as shown in embryoid body.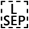Scale bar, 100 μm. J. An image of a chimeric mouse derived from si6B-mdx-ntES.

Subsequently, we evaluated whether KDM6B knockdown could improve ntES derivation. Therefore, SCNT blastocysts were derived from MERVL::tdTomato/Oct4::EGFP cumulus cells and the standard protocol was used to establish ntES. Compared with unmanipulated SCNT embryos, the efficiency of ntES derivation increased from 39.5% to 80.3% with siRNA-6B injection, and all ntES lines expressed Oct4::EGFP (Appendix Fig S6G). As SCNT can be used to consistently reprogram somatic cells to pluripotency, it is ideal for cell replacement therapies. We next used mouse tail-tip MEFs of DMD-deficient mdx mice as nuclear donors. SCNT blastocyst attachment to the feeder cell increased from 13% to 57%, and ntES derivation increased from 4% to 27% by siRNA-6B injection (Fig 6G). The si6B-mdx-ntES generated from siRNA-6B injected SCNT blastocyst showed characteristic ES morphology and expressed ES markers such as Oct4, Sox2, Ssea1, and E-Cadherin (Fig 6H). To further investigate the si6B-mdx-ntES differentiation capacity, we performed *in vitro* differentiation and *in vivo* chimera assays. si6B-mdx-ntES could efficiently give rise to three germ layer cells (Fig 6I). Furthermore, the si6B-mdx-ntES lines efficiently contributed to adult chimeric mice (Fig 6J).

### SCNT Embryonic Transcriptome Upon KDM6B Knockdown Resembled *In Vivo* Fertilized Embryos

Having demonstrated that KDM6B knockdown markedly improved SCNT efficiency, we next evaluated corresponding changes at the molecular level. We first used a qPCR assay to detect the 2-cell embryo-specific transcripts (Appendix Fig S7A). We found that *Zscan4, Gm6763, Eif1a*, and *MERVL* levels were higher in NT blastocysts with siRNA-6B injection than in the control. Interestingly, *MERVL* was strongly upregulated, but other repeat elements, such as *LINE-1* and *IAP* (intracisternal A particles), were unaffected. These results suggested that the knockdown of KDM6B improved the developmental potential of SCNT embryos by increasing ZGA-related transcripts. To further verify this result, we used single-cell RNA sequencing (scRNA-seq) to evaluate the transcriptome in siRNA-6B-injected SCNT embryos. We also noticed that injection of siRNA-6B does not make every SCNT embryos active MERVL::tdTomato expression and reach the blastocyst. We combined live-cell imaging, blastomeric biopsy, and scRNA-seq to accurately characterize the molecular characteristics (Fig 7A; Movies EV3). We first confirmed that the removal of a single blastomere at the 2-cell stage did not influence the developmental capacity (Appendix Fig S7B, C; Movies EV4). Using this system, we removed one blastomere from siRNA-6B-injected MERVL::tdTomato-SCNT 2-cell stage embryos for scRNA-seq (referred to as si6B-NT); the remaining blastomeres were monitored by live-cell imaging system for blastocyst formation and tdTomato expression. We also generated scRNA-seq profiles for normal SCNT 2-cell embryos (NT-2), and the publicly available fertilized 2-cell embryo RNA-seq dataset was harvested as WT-2 [45]. Finally, we obtained > 65 million 90-bp reads per sample, with at least 72.8% of the reads aligning to the mouse genome. Two biological replicates for each sample demonstrated high reproducibility (Appendix Fig S7D). Compared to NT-2 embryos, 1,175 genes were highly expressed in siRNA-6B-injected embryos (FC > 5, FPKM > 5; Fig 7B). We next focused on the expression of 7,773 representative ZGA-related gene [45], because we supposed that knockdown of KDM6B to promote ZGA in SCNT embryos. A pairwise comparison of the transcriptomes of NT-2, si6B-NT, and WT-2 embryos identified 1,813 differentially expressed ZGA-related genes (FC > 5, FPKM > 5), and these DEGs (differentially expressed gene) could be classified into two groups (designated Group1 and Group2) by an unsupervised hierarchical cluster analysis (Fig 7C; Dataset EV1, 2). Group2 genes were significantly more highly expressed in SCNT embryos injected with siRNA-6B than in the NT-2 embryo. To further investigate whether these DEGs cause developmental issues in SCNT embryos, we used GO (Gene ontology) and KEGG (Kyoto encyclopedia of genes and genomes) to analyze enrichment for biological processes and pathways. Group2 genes were enriched for cell cycle, methyltransferase activity, ribosome, and mitochondrion categories. These results suggest that the dysregulation of these developmentally important genes might be a cause of SCNT failure.

**Figure. 7.**
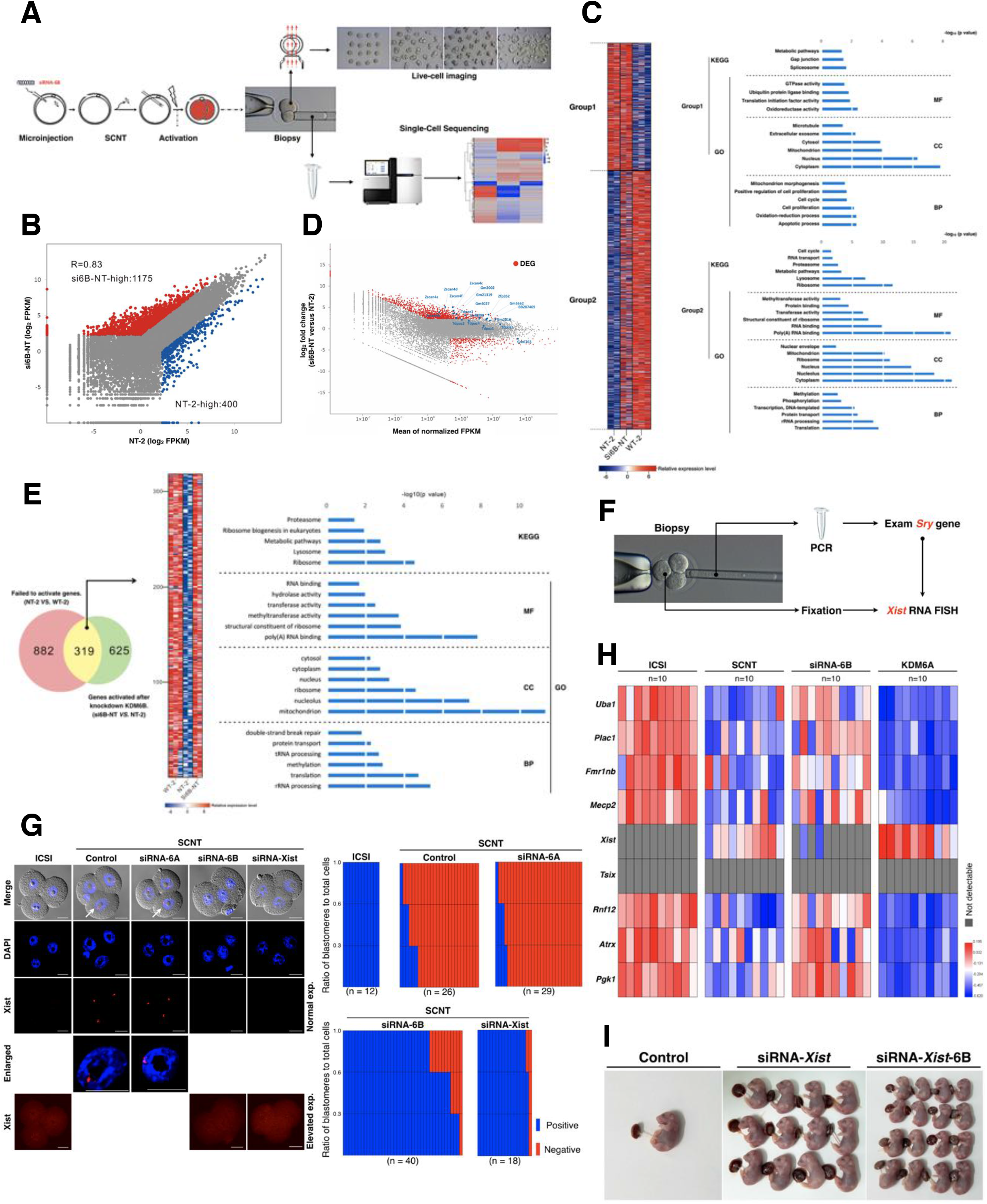
Analyses of Molecular Features of knockdown KDM6B assisted SCNT. A. Schematic illustration of the experimental procedures.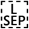We combined the live-cell imaging, blastomere biopsy and single-cell RNA sequencing to accurate analysis. B. Scatter plots comparing the si6B-NT and NT-2 genes expression. The higher expression genes in siRNA-6B and NT-2 are colored with red and blue, respectively (FC > 5, FPKM >5). C. Heatmap comparing ZGA genes expression between WT-2 and NT-2 and si6B-NT embryos (FC > 5, FPKM > 5 in each replicate; left). A total of 1,813 DEGs are classified into two groups by unsupervised hierarchical clustering. KEGG and GO analysis of the two groups by unsupervised hierarchical clustering (right). D. MA plot comparing gene expression between si6B-NT and NT-2. The data analysed derive from two independent biological replicates. Arrows represent data points outside of the plotting area. DEG, differentially expressed genes. E. Venn diagram showing the overlap between the genes that failed to be activated in SCNT 2-cell embryos and derepressed in knock-down KDM6B (left). Heatmap, KEGG and GO enrichment showing the expression pattern of 319 overlap genes (FC > 5, FPKM >5; right). F. Schematic illustration of the experimental approach. G. Representative localization of Xist expression in the nuclei of SCNT embryos injected with siRNA-6A, siRNA-6B or siRNA-Xist (left). Arrows indicate the blastomeres enlarged in the bottom panels. The ratios of blastomeres classified according to the positive or negative expression of Xist analyzed by RNA FISH (right). Each bar represents a single embryo. H. Large-scale qPCR analysis of Xist and eight X-linked genes in 4-cell stage male embryo. Each rectangle bar represents the expression value detected by single embryo RT-qPCR. The number of embryos in each group is indicated. Coloured bars indicate expression levels. I. Image of full-term cloned pups derived from NT Sertoli cells injected with siRNA-control, siRNA-Xist, or siRNA-Xist-6B.

It is well known that Zscan4 plays an important role in lengthening telomeres by recombination-based mechanisms and in maintaining genomic stability during embryonic development; the depletion of Zscan4 causes a severe delay in pre-implantation development [46]. Therefore, we further examined the DEGs between si6B-NT and NT-2 (Fig 7D; Appendix Fig S7E). We found that the knockdown of KDM6B expression increased the expression of *Zscan4* and *Eif1a-like* genes, suggesting that the knockdown of KDM6B increases the efficiency of SCNT reprogramming. Furthermore, we also identified 319 genes that were not activated in 2-cell SCNT embryos and were derepressed by KDM6B knockdown (FC > 5, FPKM > 5; Fig 7E; Dataset EV3). KEGG and GO analyses indicated these genes are enriched for methyltransferase activity, metabolic pathways, and RNA processes (Fig 7E right). Taken together, these results indicate that KDM6B knockdown can facilitate the activation of the embryonic genome in SCNT reprogramming.

### Knockdown KDM6B not Only Facilitate ZGA in SCNT, but also Impede Ectopic Xist Expression

The results above showed that 2-cell stage aberrant epigenetic reprogramming can be rescued through overexpression KDM6A or knock down KDM6B. Although aberrant SCNT-ZGA is believed to the main reason for low cloning efficiency. Another error identified in SCNT embryo is ectopic expression of the Xist (X-inactive specific transcript), which initiates X chromosome inactivation. Recently, H3K27me3 was identified as an imprinting mark for Xist [47], which prompted us to ask whether it is also responsible for fine-tuning KDM6A/B improved development of SCNT embryo. Due to exact adjustment by siRNA is technically difficult, we next primarily focused on male SCNT embryos with only a single X chromosome and never expressed at 4-cell stage. According to a previous report [48], sex screening of early mouse embryos was determined by PCR using a single blastomere biopsy at the 4-cell stage (Fig 7F). To determine whether loss of H3K27me3 modification can induce Xist derepression in embryos, we first injected KDM6A and KDM6B mRNA into ICSI derived embryos. As Inoue A et al. report [47], RNA fluorescent in situ hybridization (FISH) analysis confirmed that KDM6A/B mRNA injection induce ectopic expression of Xist, and only KDM6A in a concentration-dependent manner (Appendix Fig S7F). To evaluate the effect of fine-tuning KDM6A/B on Xist expression in SCNT embryos, we harvested Sertoli cell derived SCNT embryos for Xist RNA detection via FISH assay. As shown in Fig 7G, the majority of SCNT derived blastomere showed Xist RNA signal, and ICSI derived embryos showed no Xist signal. As expected, KDM6B knockdown by siRNA-6B led to Xist down-regulation and loss of Xist signal within the nucleus of SCNT embryos. In contrast, most of the siRNA-6A injected SCNT embryos still showed one strong Xist signal in blastomeres. Previous studies demonstrated that ectopic expression of Xist will lead to large-scale downregulation of X chromosome-linked genes in the SCNT embryos [3, 49]. The effect of siRNA-6B on ectopic Xist expression was further examined the expression levels of Xist and X-linked genes (*Tsix, Rnf12, Pgk1, Fmr1nb, Atrx, Uba1, Mecp2* and *Plac1*) via single embryo RT-qPCR (Fig 7H). Consistent with the FISH results, the significant down-regulation of Xist observed in SCNT embryos that had been injected with siRNA-6B. In contrast, Xist was significantly up-regulated in KDM6A mRNA injected SCNT embryos, and the X-linked genes were also up-regulated in siRNA-6B injected embryos.

Related studies have demonstrated that the ectopic expression of Xist in SCNT derived embryos could be corrected by siRNA-Xist, leading to more than a 10-fold increase in the birth rate of male clones [3, 49, 50]. To examine whether the combination of siRNA-Xist and siRNA-6B could further improve SCNT embryonic full-term development. We then performed embryo transfer experiments to assess the full-term developmental ability of siRNA-Xist-6B coinjected SCNT embryos. Similar to previous report [3], injected with siRNA-Xist alone improved the birth rate from 1.3% (1/77) to 11.7% (12/103) (Fig 7I; Appendix Table S4). Importantly, siRNA-Xist-6B coinjected further increased the SCNT birth rate to 21.1% (16/76). This result indicates that siRNA-6B and siRNA-Xist exert a synergistic effect on the SCNT reprogramming. Thus, knock down KDM6B not only facilitate the cloned embryos ZGA, but it can also impede ectopic Xist expression in SCNT reprogramming.

## Discussion

Pre-implantation embryogenesis encompasses several critical events, especially the activation of ZGA-related genes. In 2014, Matoba and colleagues identified reprogramming resistant regions (RRRs), which are enriched for the histone modification H3K9me3 [7]. Liu and colleagues proved that excessive H3K9me3 modifications would lead to ZGA failure [24]. Therefore, ZGA is indispensable for somatic cell reprogramming [26]. It is noteworthy that a lack of relevant animal models has hampered precise spatiotemporal detection and critical evaluations of the efficacy of ZGA in SCNT reprogramming. Immunocytochemistry requires sample fixation and is insufficient for real-time monitoring of ZGA events. To the best of our knowledge, the present study generated the first MERVL::tdTomato transgenic mice. To detect the efficiency of SCNT reprogramming, we crossed the MERVL::tdTomato mouse strain with the Oct4::EGFP transgenic mouse strain (also known as OG2) [51]. The compound homozygous MERVL::tdTomato/Oct4::EGFP double transgenic mice provide the opportunity for serial real-time monitoring of ZGA and reprogramming efficiency.

The MERVL::tdTomato/Oct4::EGFP SCNT-embryos can be divided into three groups: MERVL^−^/Oct4^−^, MERVL^+^/Oct4^−^, and MERVL^+^/Oct4^+^. Only a small proportion of reconstructed embryos were labeled by both reporters (MERVL^+^/Oct4^+^), and we never found MERVL^−^ /Oct4^+^ SCNT embryo. We only detect moderate H3K27me3 modifications in the MERVL^+^ SCNT- and ICSI-embryos at the 2-cell stage, but we clearly detected strong H3K27me3 staining in the MERVL^−^ SCNT embryos. Although the H3K27me3 defect in SCNT embryos has been observed, previous studies have reported the loss of H3K27me3 in ICM cells of most SCNT embryos [52]. This difference might be explained by a difference in the time of embryo collection between studies. Our scRNA-seq transcriptome also demonstrated that ZGA-related genes failed to be properly activated in MERVL^−^ SCNT compared with *in vitro* fertilization embryos. Furthermore, a high H3K27me3 level is detrimental to bovine SCNT embryonic development, consistent with porcine SCNT reprogramming [53]. Collectively, these results suggested that SCNT embryos are H3K27me3-defective at the ZGA stage, which serves as another barrier to mouse, bovine, and porcine SCNT reprogramming.

In a series of rigorous experiments, we demonstrated that only injection with a low concentration (20 or 50 ng/μl) of KDM6A mRNA could facilitate the cloned embryos ZGA, and improve the pre-implantation developmental potential. Although we were able to efficiently obtain SCNT blastocysts by KDM6A injection, but failed to obtain live cloned pups. In recently, one study claimed that injection with a higher concentration (1,000 ng/μl) of KDM6A mRNA can improve the SCNT embryo preimplantation development [54], which were contrary to present study. Furthermore, Bai et al. only found high efficiency in preimplantation development of SCNT embryos by reducing H3K27me3, but whether the post-implantation development of SCNT embryos can also be improved is not test. In addition to abnormal ZGA, another SCNT reprogramming obstacle is aberrant Xist activation following SCNT [55]. The downregulation of X-linked genes is mainly caused by the ectopic expression the Xist, which responsible for the X chromosome inactivation (XCI). Deletion of Xist or repression of Xist expression by siRNA can elevate about 10-fold normal birth rate of mouse cloning [3, 49]. Bai et al. claimed that H3K27me3 removal corrected SCNT-specific aberrant XCI status in cloned embryos. This was especially of interest since it was recently reported that H3K27me3 serves as the imprinting mark of Xist, and loss of H3K27me3 induces Xist ectopic expression [47, 50]. To further determine the role of KDM6A in the XCI of SCNT, we injected KDM6A mRNA (1,000 ng/μl) into SCNT embryos, and found that the developmental efficiency of SCNT embryos was reduced, while many X-linked genes were consistently repressed. In contrast, knockdown of KDM6B could increase the SCNT embryo birth rate as well as the efficiency of DMD-specific NTES derivation. Thus, knock down KDM6B not only facilitate the cloned embryos ZGA, but it can also impede ectopic Xist expression in SCNT reprogramming.

Previous studies have shown that the knockdown of KDM6B or over-expression of KDM6A in MEFs results in significantly more iPSC colonies compared with wild-type cells [56, 57]. Interestingly, the knockdown of KDM6B in SCNT embryos leads to a moderate increase in the expression of KDM6A, consistent with our previous findings in mouse parthenogenetic embryos [42]. While our paper was under preparation, another study reported the identification of H3K27me3-dependent imprinting genes (which include *Gab1, Sfmbt2* and *Slc38a4)*, and previous studies have shown that these genes exhibit a loss of imprinting in SCNT embryos [58, 59]. This provides an explanation for why the knockdown of the H3K27me3 demethylase KDM6B promotes SCNT efficiency. In addition to the silencing of the histone modification, a recent study found that H3K4me3, an activating modification, is also obstacle to reprogramming [29]. The findings of the present study in combination with previous results in the field indicate that there are many obstacles in SCNT reprogramming. Further studies should focus on identifying the core obstacle.

## Materials and Methods

### Ethics statement

All studies adhered to procedures consistent with the National Research Council Guide for the Care and Use of Laboratory Animals and were approved by the Institutional Animal Care and Use Committee at Inner Mongolia University.

### Animals

C57BL/6N, DBA/2 and BDF1 (C57BL/6N × DBA/2) F1 strains of mice were purchased from Vital River Laboratories (China). Pseudopregnant CD1 or Kun-Ming (KM) white mice were used as embryo recipients. In order to detect reprogramming by means of Oct4 promoter driven EGFP, BDF1 mice were replaced with OG2 mice that carry an Oct4-EGFP transgene (JAX stock number 004654). All the MERLV::tdTomato transgenic mice are syngeneic and bred by the same positive Founder (F0). All the embryos used in the experiment were produced by MERVL::tdTomato sperm and MII oocytes from the littermates of transgenic mice. The copy numbers of MERLV::tdTomato were detected by previously reported methods [60, 61]. In brief, we detected approximately 200 copies of MERLV::tdTomato in reporter transgenic mice as determined by quantitative PCR. Furthermore, the MERLV::tdTomato transgene copy number was stable throughout the F20 generations.

### Superovulation and *in vivo* fertilization

Chemicals were purchased from Sigma Chemical Co. (USA) unless otherwise indicated. Superovulation was done as previously described [42]. Briefly, BDF1 female (6 ~ 8 weeks old) mice were superovulated by intraperitoneal injection of pregnant mare serum gonadotropin (PMSG; Sansheng, China, 10 IU) and human chorionic gonadotropin (hCG; Sansheng, China, 10 IU) 48 h apart. Mice were sacriced by cervical dislocation and cumulus-oocyte complexes (COCs) were collected from oviducts 14 h post hCG. For zygotes, superovulation of 7-to 8-week-old BDF1 females mated with males of the same strain. Successful mating was confirmed by the presence of vaginal plugs. The cumulus cells were dispersed by 0.3 mg/mL hyaluronidase in M2 medium (Millipore, USA).

### *In vitro* mRNA synthesis, siRNA construction and microinjection in oocytes

The coding region of KKDM6A and KDM6B was amplified from mouse tail tip genome. Forward and reverse primers contained T7 promoter and HA sequences, respectively. To prepare mRNAs for microinjection, pT7-Cas9 (OriGene, GE100014), KKDM6A and KDM6B expression vectors were linearized and subjected to phenol-chloroform extraction and ethanol precipitation. The mRNA synthesized with the mMESSAGE-mMACHINE T7 Ultra Kit (Thermo, USA) according to the manufacturer’s instructions. Two different siRNA species targeting KDM6B were designed and synthesized using the silencer siRNA construction kit (Ambion, USA) following the manufacturer’s instructions. A commercially available siRNA without any specificity to known genes was used as control. As previously described [42], with minor modifications, 8 pL of siRNA-6B or siRNA-control was microinjected into the cytoplasm of denuded MII oocytes. Oocytes were injected using Piezo-operated blunt-end micropipette (3 ~ 5 μm internal diameter). After injection, oocytes were kept at RT for 30 min and then moved into the incubator.

### Transgenic mice generation

The MERVL::tdTomato vector was a gift from Samuel Pfaff (Addgene 40281). The vector was linearized with the enzyme. The pronuclear microinjection for the production of transgenic mice followed previously published studies [62]. Briefly, the linearized vector was injected into the well-recognized pronuclei, in M2 medium. Injected zygotes were transferred into pseudopregnant female mice (~30 zygotes per mouse) after 4 h recovery culture in KSOM-AA medium. For founder identification, Tail tips were subjected to standard DNA-extraction procedures. For identification MERVL::tdTomato of founders, the extracted DNA was amplified with MERVL::tdTomato primers flanking the target sites (Appendix Table S5). Primers were synthesized by Takara Biotechnology Dalian Co. Ltd (Dalian, China). The amplified DNA fragments were subjected to TA cloning and sequencing. The founder mice were crossed to the littermates of founder mice for four generations to produce homozygous MERVL::tdTomato mice. We intercrossed MERVL::tdTomato mice with homozygous Oct4::EGFP transgenic mice (OG2) for six generations to produce MERVL::tdTomato/Oct4::EGFP dual reporter mice.

### SCNT, ICSI, and IVF

The mouse-SCNT, was done as previously described [63]. Briefly, MII oocytes after a brief culture in KSOM-AA medium, groups of ~50 oocytes were transferred to a chamber containing oil-covered M2 supplemented with 5 μg/mL cytochalasin B (CB). The spindle chromosome complex (SCC) was removed by a blunt Piezo-driven pipette (PrimeTech, Japan) on a 37 °C heating stage of an inverted microscope (Nikon, Japan). The nuclei of donor cumulus cells, Sertoli cells, or C57-MEF cells, a small cell (< 10 μm) was drawn in and out of the injection pipette until its plasma membrane was broken and was then injected into enucleated oocytes. For the mdx-MEF cells, live cells with a diameter of 10~15 μm were selected. The reconstructed embryos were cultured in M199 medium (Thermo, USA) containing 10% fetal calf serum (FCS; Hyclone, USA) for 1~3 h before activation treatment. The reconstructed embryos were activated in Ca^2+^ free KSOM medium containing 10 mM strontium and 5 μg/mL CB for 6 h. Activated constructs were thoroughly washed and cultured in G1 and G2 medium (Vitrolife, Sweden). The bovine-SCNT, bovine oocytes obtained by aspirating follicles on slaughterhouse-derived ovaries. We cultured immature cumulus-oocyte complexes in M199 medium supplemented with 10% FCS, 0.2 mM pyruvate, 200 μg/mL gentamicin, 0.5 mg/mL luteinizing hormone and 1 mg/mL estradiol for 16 to 18 h at 38.5 °C with 5% CO_2_ in the air. After 18 h the start of maturation, cumulus cells were removed from the oocytes, and oocytes with extruded first polar bodies were selected as MII oocyte. Oocytes enucleated using a beveled glass pipette by aspirating the first polar body and the MII plate in a small amount of surrounding cytoplasm in M199-HEPES medium containing 5 μg/mL CB. In some experiments, we labeled oocytes with DNA fluorochrome (Hoechst 33342) before enucleation; to ensure removal of the oocyte chromatin, we exposed the aspirated cytoplasm to UV light to examine the enucleation. The donor cells were injected into the perivitelline space of each enucleated oocytes by using the same slit in the zona pellucida as made during enucleation. Then, we fused nuclear transfer couplets in sorbitol fusion medium by applying a single electric pulse (1.2 kV/cm for 30 μs). One hour after fusion, the fused embryos using 5 μM ionomycin for 5 min, followed by five hours of treatment with 10 μg/mL cycloheximide (CHX). The reconstructed embryos were cultured and allowed to develop *in vitro* up to the 8-cll or blastocyst stage. The mouse-ICSI, MII oocytes were collected 18 h post hCG. The sperm were collected from epididymis of 9-week-old BDF1 mice and washed in M2 medium, then suspended in M2 medium supplemented with 10% polyvinylpyrrolidone (PVP). The MII oocytes were placed in a drop of M2 medium and one sperm head was injected into a MII oocyte by piezo-micromanipulator. The surviving embryos were collected and cultured in G1 and G2 medium. The ICSI was done as previously described [64]. Only the sperm head was injected into the oocyte. After 30 min of recovery, the ICSI-generated embryos were washed several times and cultured in KSOM-AA medium at 37 °C in a 5% CO2 in air atmosphere. The mouse-IVF, sperm was obtained from the cauda epididymis of male mouse and incubated at 37 °C for 1 h in HTF supplemented with 5% FBS before the addition of the COCs. The presence of pronuclei was scored 6 h after the initiation of the IVF reaction. After gamete coincubation, the zygotes were collected and cultured in G1 and G2 medium. The bovine-IVF, COCs matured for 24 h were co-incubated with sperm (10^6^ spermatozoa/mL; thawing semen in 37 °C water) in IVF medium at 38 °C in 5% CO_2_ in air for 20 hours. The IVF medium consisted of NaCl (114 mM), KCl (3.15 mM), NaH_2_PO_4_ (0.39 mM), Na-Lactate (13.3 mM), CaCl_2_ (2 mM), MgCl_2_ (0.5 mM), Na-Pyrovate (0.2 mM), Penicillin (50 IU/mL), Streptomycin (50 μg/mL), NaHCO_3_ (25 mM), Heparin (10 μg/mL), Penicillamine (20 μM), Hypotaurine (10 μM), Epinephrine (1 μM), bovine serum albumin (BSA; 6 mg/mL). Presumptive zygotes were vortexed for 2 min to separate cumulus cells. Groups of ~40 presumptive zygotes were cultured in 500 μL drops of SOF medium under mineral oil at 38.5 °C, 5% CO_2_ in humidified air. 72 h after insemination, 5% FCS was added to the culture media.

### ntESCs derivation, chimeric mice and embryo transfer

Blastocysts were denuded by Acidic Tyrode’s solution and plated on mitomycin treated MEF feeder layers in a 96-well plate. The ntES cells derivation medium contains Knockout-DMEM (Thermo, USA) supplemented with 15% (v/v) knockout-serum-replacement (KSR; Thermo, USA), 1mM GlutaMAX (Thermo, USA), 0.1 mM mercaptoethanol, 1% nonessential amino acid (Thermo, USA), penicillin/streptomycin (100×; Thermo, USA), nucleosides (100×; Thermo, USA) and 1,000 U/mL LIF (Thermo, USA). The ntESC colonies formed with culturing for 10 days, and were picked and transferred for cell passage. The expansion of ES cells was performed by routine culture. For chimeric experiments, ntESCs were used one day before passaging, which showed an optimal undifferentiated morphology. The ntESCs were microinjected into CD1/KM blastocysts using a piezo-microinjection pipette. After culturing for 3 h, the embryos were transplanted into the uterus of pseudo-pregnant mice (~20 mbryos per mouse). The 2-cell stage SCNT, siRNA-6B, or KDM6A/B injected embryos were transferred to the oviducts of E0.5 pseudo-pregnant (~20 mbryos per mouse). The embryos were recovered by caesarian section on the E8.5, E14.5, or E19.5. The cloned pups nursed by lactating CD1/KM females. SSLP analysis was performed for D6Mit15, D2Mit102, D11Mit236, D4Mit204 and EGFP. The primer information is presented in Appendix Table S5.

### Immunofluorescence staining and quantification analysis

Embryos and ntESCs were rinsed three times in phosphate buffered saline (PBS) with 0.3% BSA, fixed with 4% paraformaldehyde (PFA) overnight at 4 °C and then permeabilized with 0.2% (*vol./vol*.) Triton X-100 for 15 min at room temperature, followed by by washing thoroughly in PBS containing 0.3% BSA. Fixed samples were blocked in 0.05% Twesen-20 in PBS containing 3% BSA (PBST) at 37 °C for 1 h and then incubated with the primary antibodies overnight at 4 °C. After blocking and simultaneous incubating with primary antibodies: anti-H3K27me3 (Millipore, ABE44, USA), anti-H3K27me2 (Abcam, ab24684, USA), anti-H3K4me3 (Abcam, ab213224, USA), anti-H3K9me3 (Abcam, ab176916, USA), anti-HA (Santa Cruz, sc-7392, USA), anti-MuERVL-Gag (Epigentek, A-2801-100, USA), anti-Oct4 (Santa Cruz, sc-8629, USA), anti-Sox2 (Santa Cruz, sc-17319, USA), anti-Cdx2 (Abcam, ab76541, USA), anti-Nanog (Abcam, ab107156, USA), anti-Ssea1 (Santa Cruz, sc-21702,USA), anti-E-cadherin (Abcam, ab40772,USA), anti-Nestin (Santa Cruz, sc-21247, USA), anti-Brachyury (Santa Cruz, sc-17745, USA), anti-Gata4 (Santa Cruz, sc-1237, USA). After incubating, the samples were needed to wash several times in PBST and then incubated with appropriate secondary antibodies conjugated with Alexa Fluor 594 and Alexa Fluor 488 (Thermo, USA) for 1 h at 37 °C. For imaging the embryos were mounted in 10 μl anti-fade solution with DAPI (Thermo, USA) and compressed with a coverslip. After mounted on glass slides and examined with a confocal laser-scanning microscope (Nikon, Japan). For fluorescence quantification, the signal intensity was analyzed as described previously [65–67] (PMID: 18784248, 20422712, 25925669). Briefly, nuclei of blastomeres were identified by DAPI staining. Quantification analysis of fluorescence intensity in nuclei or cytoplasmic areas was performed using ImageJ software (NIH, Bethesda, MD, USA; http://rsbweb.nih.gov/ij/). In addition, at least three different cytoplasmic areas were delineated for normalization to background. The average pixel intensity of the nuclear areas was calculated by ImageJ, and then normalized by dividing by the average pixel intensity of the background areas.

### RNA-FISH

RNA-FISH on preimplantation embryos was performed as previously described [47, 68]. Briefly, the embryos were fixed in 2% PFA in PBS containing 0.5% Triton X-100 for 20 min at room temperature. After three washes with 0.1% PVP/PBS, embryos were treated with 0.1 N HCl containing 0.02% Triton X-100 for 15 min at 4 °C. After three washes with 0.1% PVP/2× SSC, embryos were incubated in a series of 10%, 20%, and 50% formamide/2× SSC. The samples were covered with mineral oil, heated for 30 min at 80 °C, and then incubated for ~30 min at 37 °C. Next, the fixed embryos were performed using ViewRNA ISH Cell Assay Kit (Thermo, USA) based on the manufacturer’s instructions. Custom-designed ViewRNA Cell Plus Probe against Xist (Thermo, VX-06, USA). The embryos were then counterstained with DAPI, and fluorescence was detected under a laser-scanning confocal microscope (A1+, Nikon, Japan). Gender identification was performed by PCR according to the methods described above.

### RNA extraction and RT-qPCR

As previously described [42], total RNA was extracted using the Pico-Pure RNA Isolation Kit (Thermo, USA) according to the manufacturer’s instructions. Total RNA was extracted from each pool of embryos (n = 3 pools of 20 oocytes or embryos per time point), and residual genomic DNA was removed by DNase I digestion, using an RNase-Free DNase kit (Qiagen, Germany). Reverse transcription was performed using SuperScript III (Thermo, USA) following the manufacturer’s instructions. Quantitative RT-PCR was performed using a SYBR-Taq Master Mix (Applied-BioSystems, USA) and signals were detected with ABI7500 real-time PCR System (Applied-BioSystems, USA). Analysis of relative gene expression was measured using the 2^(-Delta Delta Ct) method. For the single embryo RT-qPCR, was done as previously described [69]. Briefly, embryonic total RNA was extracted using an RNeasy Micro Kit (Qiagen, Germany) and treated with DNase following the manufacturer’s instructions. mRNAs were reverse by SuperScriptIII Reverse Transcriptase kit (Thermo, USA). For quantitative gene expression analysis with high specificity, TaqMan probes (Thermo, USA) were used in single embryo RT-qPCR assays, and the expression levels of all embryos were normalized to the average expression levels of ICSI group. All the TaqMan probes and primer sets used in this study are shown in Appendix Table S5.

### Embryo biopsy, library construction and single-cell RNA-seq

The 2-/4-cell embryos were transferred into Ca^2+^ and Mg^2+^ free KSOM-AA medium for 1 h to disrupt cell adhesion, and were then transferred to Ca^2+^ and Mg^2+^ free M2 medium on the micromanipulation dish. The zona pellucida was penetrated by a blunt Piezo-driven micropipette (~ 30 μm inner diameter) and one blastomere was gentle aspirated from each manipulated embryo, the rest blastomere were cultured in G1+G2 (1:1) medium with 5% CO_2_ at 37 °C. Control (nonbiopsied) embryos were from the same SCNT cohorts and were cultured under the same conditions as their biopsied counterparts but were not micromanipulated. The isolated single blastomere was washed twice in PBS-BSA (0.1%) and hold individual blastomeres before placing in lysis buffer and stored in liquid nitrogen. The single-cell RNA-seq method followed previously published studies [24], only capture mRNAs with a poly(A) tail. Library construction was performed following the Illumina manufacturer’s instructions and sequencing was performed at the BGI (China). Paired-end sequencing was further performed on the Illumina Hiseq2000 platform. The sequencing reads that low quality and adapters were pre-filtered before mapping. Filtered reads were mapped to the mm9 genome using Tophat (v1.3.3) with default parameters, and evaluated using RseQC (v2.3.4). Transcriptional profiling was done as described [24]. Briefly, data normalization was performed by transforming uniquely mapped transcript reads. Genes with low expression in all stages were filtered out, and quantified to FPKM (fragments per kilobase of exon model per million mapped reads) using Cufflinks (v1.2.0) to eliminate the effects of sequencing depth and transcript length. Some analyses were performed using R software.

### Live-cell Imaging procedures

Live-cell Imaging was done as previously described [42, 70]. Briefly, the embryos were transferred to drops of KSOM-AA medium, and placed in the incubator (Tokai Hit, Japan) on the microscope stage (A1+, Nikon, Japan) and incubated at 37 °C under 5% CO_2_ in air, Images were acquired by an electron multiplying charge-coupled device (EM-CCD) camera (iXon 897, Andor Technology, UK). Images were taken over 96 h at 10 or15 min intervals. Live-cell Imaging system was housed in a dark room at 27 °C.

### Data availability

Sequencing data have been deposited in the NCBI sequence read archive (SRA) under accession code SRR6024636. All other data are available from the authors upon reasonable request.

## Acknowledgements

We thank our laboratory colleagues for their assistance with the experiments; Shaorong Gao (Tongji University), Zhiming Han (Chinese Academy of Sciences), and Qing Xia (Peking University) for technical assistance. We also thank Yuan Li (New York University) for critical reading and editing of this manuscript; Yi Zhang (Harvard Medical School) for sharing the list of RRRs. This study was supported by the National Transgenic Animal Program (2016ZX08007-002) and Basic Research Project in the Inner Mongolia Autonomous Region (201503001).

## Author contributions

L.Y. and G.L. conceived and designed the study. L.Y., L.S., and X.L. performed the experiments; L.Y., L.S., X.L, L.B., and G.L. analyzed the data. L.Y. and G.L. supervised the project. L.Y. and G.L. wrote the manuscript.

